# Structural basis for EROS binding to human phagocyte NADPH oxidase NOX2

**DOI:** 10.1101/2023.09.11.557130

**Authors:** Shiyu Liang, Aijun Liu, Yezhou Liu, Fuxing Wang, Youli Zhou, Tao Wang, Zheng Liu, Ruobing Ren, Richard D. Ye

## Abstract

EROS (essential for reactive oxygen species) is a recently identified molecular chaperone of NOX2 (gp91^phox^), the catalytic subunit of phagocyte NADPH oxidase. Deficiency in NOX2 expression or function due to genetic mutations leads to chronic granulomatous disease (CGD) with recurrent bacterial and fungal infections. To delineate how EROS interacts with NOX2, we solved the cryo-EM structure of the EROS-NOX2-p22^phox^ heterotrimeric complex. EROS binds to NOX2 in plasma membrane through its anti-parallel α-helices H1 and H2, and in cytoplasm through multiple β-strands that form hydrogen bonds with the C terminal fragment of NOX2. EROS binding alters the conformation of the TM2 and TM6 transmembrane helices, increases the distance between the two hemes, and causes dislocation of the binding site for flavin adenine dinucleotide (FAD). EROS colocalizes with NOX2 on cell surface of neutrophil-like HL-60 cells and forms a heterotrimer with mature NOX2-p22^phox^ in transfected cells. Phorbol myristate acetate, an activator of NOX2, induces dissociation of EROS from NOX2 in a NanoLuc complementation assay with concurrent production of superoxide in reconstituted cells. Taken together, these findings provide a structural basis for EROS-NOX2 interaction and suggest a previously unidentified function of EROS in regulating NOX2 activation.

## INTRODUCTION

Chronic granulomatous disease (CGD) is a genetic disorder characterized by frequent and sometimes life-threatening bacterial and fungal infections [1]. Children with CGD are often healthy at birth, but develop symptoms of severe infections in infancy and early childhood [2]. Phagocytes isolated from CGD patients are found defective in bactericidal functions but normal in phagocytosis, leading to formation of granulomas that isolate the engulfed bacteria and fungi [3, 4]. Genetic studies of CGD patients led to the finding of mutations in NADPH oxidase composed of the integral membrane proteins gp91^phox^ (NOX2) and p22^phox^, the cytosolic factors p67^phox^, p47^phox^, p40^phox^, and the small GTPase Rac [5, 6]. In resting phagocytes, NOX2 is inactive and the cytosolic factors are separate from the membrane components. The activation of NADPH oxidase requires translocation of the cytosolic factors to plasma or phagosome membrane and assembly of an active complex for one-electron reduction of molecular oxygen and superoxide production by the catalytic subunit [6–8]. In phagosomes, superoxide is rapidly converted to other reactive oxygen species (ROS) such as hydrogen peroxide (H_2_O_2_) and hypochlorous acid, effectively killing the engulfed microorganisms [8]. Genetic mutations in NOX2 may lead to defective catalytic activity of the oxidase in X-linked inheritance, whereas mutations in other components of phagocyte NADPH oxidase may lead to autosomal recessive CGD [1, 2]. A search of the human genome identified homologues of the gp91^phox^-encoding genes and the establishment of the NADPH oxidases (NOX) family [9]. Of the 7 NOX members, NOX2 (gp91^phox^) is the first identified and most extensively studied catalytic subunit of this oxidase family [10], largely due to a close relationship between CGD and genetic defects in NOX2.

EROS (<u>E</u>ssential for <u>R</u>eactive <u>O</u>xygen <u>S</u>pecies) was identified in 2017 during a study of a CGD case [11]. Subsequent studies have confirmed that genetic defect of EROS can result in CGD primarily due to failure in gp91^phox^ expression [12–14]. EROS, that contains 187 amino acids, was predicted to be a membrane protein with two α-helices. The gene that encodes EROS was then given the symbol *CYBC1* [11]. It was subsequently confirmed that EROS associates closely with NOX2, in particular the newly synthesized NOX2 polypeptide of 58 kDa [14]. An endoplasmic reticulum (ER) protein, EROS binding of NOX2 protects the catalytic subunit from rapid degradation [14]. Published work has shown that EROS binding precedes heme incorporation into and p22^phox^ association with NOX2 [14]. Based on these findings, EROS is defined as a molecular chaperone of NOX2 essential for its maturation including transport through the Golgi apparatus and acquisition of additional carbohydrate moieties on top of its N-linked high-mannose glycans. This function of EROS has been confirmed in genetically altered mice and in transfected cell lines [14], suggesting a critical role for EROS in NOX2 expression and transport to plasma membranes.

Biochemical studies have shown that EROS directly binds NOX2, but the domains and amino acid residues involved in this association have not been identified. Moreover, it is unclear whether EROS continues to bind NOX2 as it matures and whether this association has a functional impact on NOX2 activation. To address these questions, we characterized cell surface expression of EROS in neutrophil-like HL-60 promyelocytic cells and found EROS co-localization with NOX2. Using recombinant human EROS, we reconstituted a protein complex containing EROS, NOX2 and p22^phox^. An analysis of samples collected from size-exclusion chromatography found both mature NOX2 and EROS in the same fraction. We then resolved the structure of the protein complex by cryogenic electron microscopy (cryo-EM). Our structural model showed key interacting residues between the transmembrane helices of EROS and NOX2 in its transmembrane domains (TM) 2 and TM6. There are extensive interactions between the C terminal fragments of EROS and NOX2 that effectively prevents access of flavin adenine dinucleotide (FAD) and NADPH to the dehydrogenase (DH) domain of NOX2. These findings indicate that EROS binding to NOX2 stabilizes the catalytic subunit in an inactive conformation.

## RESULTS

### EROS binds mature NOX2 and colocalizes with NOX2 on plasma membrane

Recent studies have shown that EROS binds NOX2 in endoplasmic reticulum (ER), an event that occurs early in NOX2 biosynthesis [13, 14]. EROS associates with NOX2 prior to heme incorporation and formation of the NOX2-p22^phox^ heterodimer, when NOX2 has acquired N-linked high-mannose core [14]. In the ER, EROS serves as a molecular chaperone for NOX2 maturation and prevents degradation of newly synthesized NOX2 (58 kDa) polypeptide [14]. However, it is unclear whether EROS remains associated with NOX2 after its maturation, and if EROS binding affects the activation of NOX2. We have analyzed cell surface expression of EROS in differentiated neutrophil-like HL-60 cells (dHL-60). As shown in Fig. 1A, EROS (green fluorescence) appeared as halos around the dHL-60 cells, and colocalized with NOX2 (red fluorescence). Colocalization of EROS and NOX2 was also observed in COS-7 cells cotransfected with the expression constructs of EROS and NOX2 (Fig. 1B), but their plasma membrane localization was uncertain. To confirm that EROS and NOX2 were colocalized on cell surface, dHL-60 cells were subject to flow cytometry using a FITC-labeled anti-EROS and anti-NOX2 (7D5) [15] followed by addition of PE-labeled anti-mouse secondary antibody. As shown in Fig. 1C, 77% of the tested cells showed double-positive staining indicating colocalization of the two membrane proteins on the surface of these cells. To further determine if EROS associates with mature NOX2, expression vectors of EROS, NOX2 and FLAG-tagged p22^phox^ were cotransfected into Expi-HEK293F cells. Forty-eight hours after transfection, the cells were collected and total cell lysates were prepared. After affinity purification with anti-FLAG, the eluent was incubated with the antibody fragment 7G5 Fab that recognizes an extracellular loop of mature NOX2 [16]. Size-exclusion chromatography (SEC) was performed and two major peaks were obtained (Fig. 1D). The collected peak fractions were resolved on SDS-PAGE. As shown in Fig. 1E), three species were identified from the large peak fraction, containing respectively the mature NOX2 (approx. 91 kDa), 7G5 Fab (approx. 26 kDa), and EROS and p22^phox^ running at approx. 22 kDa. Taken together, these results provide strong evidence that EROS associates with mature NOX2 on cell surface.

**Figure 1.**
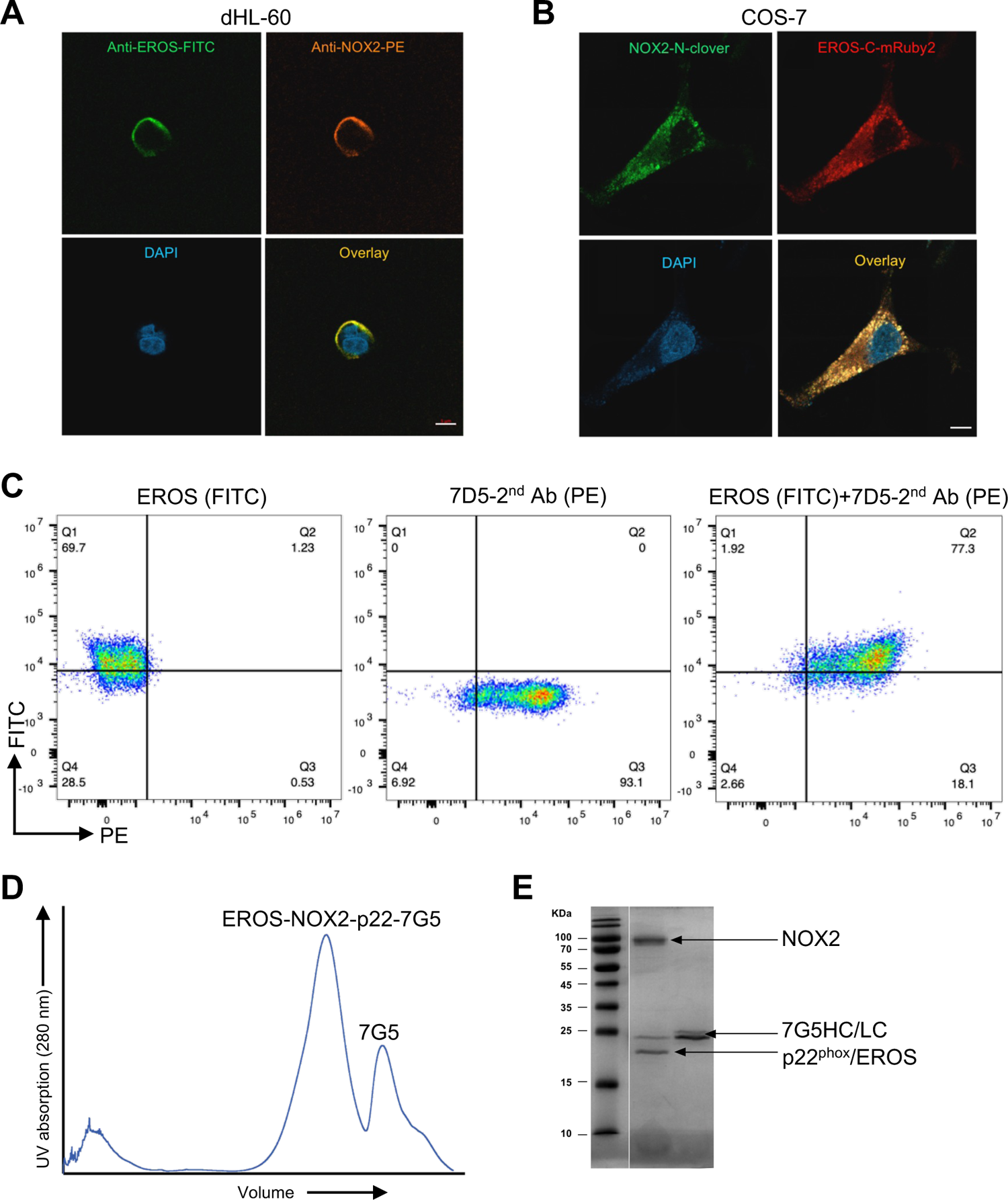
Colocalization of EROS with NOX2 on cell surface and interaction of EROS with mature NOX2. **A.** Images from confocal fluorescent microscopy showing the expression of EROS (FITC, green fluorescence) and NOX2 (PE, red fluorescence) on the surface of differentiated HL-60 cells (dHL-60). Colocalization of the two membrane proteins (yellow, merged image) was shown in the lower right panel. Scale bar: 5 μm. **B.** Colocalization of EROS and NOX2 in transiently transfected COS-7 cells. The cells were cotransfected with NOX2-N-Clover (green) and EROS-C-mRubby2 (red). Confocal microscopy images were taken 24 h after transfection. **C.** Verification of cell surface expression of EROS and NOX2 in dHL-60 by flow cytometry, using the primary antibodies anti-EROS-FITC and anti-NOX2 (7D5) plus PE-labeled goat anti-mouse secondary antibody for 1-h incubation. **D.** Size-exclusion chromatography of the EROS-NOX2-p22^phox^-7G5 Fab complex on Superose 6. The two major peaks were collected and subjected to SDS-PAGE. **E.** NOX2 was detected at an expected molecular weight of ∼91 kDa, indicating that the EROS-associated NOX2 is in mature form.

### Cryo-EM structure of the EROS-NOX2-p22^phox^ complex

The large peak fraction from SEC contained an EROS-NOX2-p22^phox^ complex stabilized by the 7G5 Fab. This fraction was subject to structural analysis by cryo-EM as detailed in *<u>Methods</u>*. Electron density map of the complex (Fig. 2A) showed an overall resolution of 3.56 Å. The overall architecture of the NOX2-p22^phox^ complex, including the 6 transmembrane domains (TM) of NOX2 and the 4 TM bundles of p22^phox^, were essentially the same as reported recently [16, 17]. EROS in this complex is associated with TM2 and TM6 of NOX2 through the two α helices H1 and H2 (Fig. 2A). An intracellular view of the complex (right panel in Fig. 2A) found that EROS and p22^phox^ are on opposite sides of NOX2 with a plane angle of 177°. With EROS included, the complex has a maximal base width of 85Å, increasing from 60Å in the absence of EROS [17].

**Figure 2.**
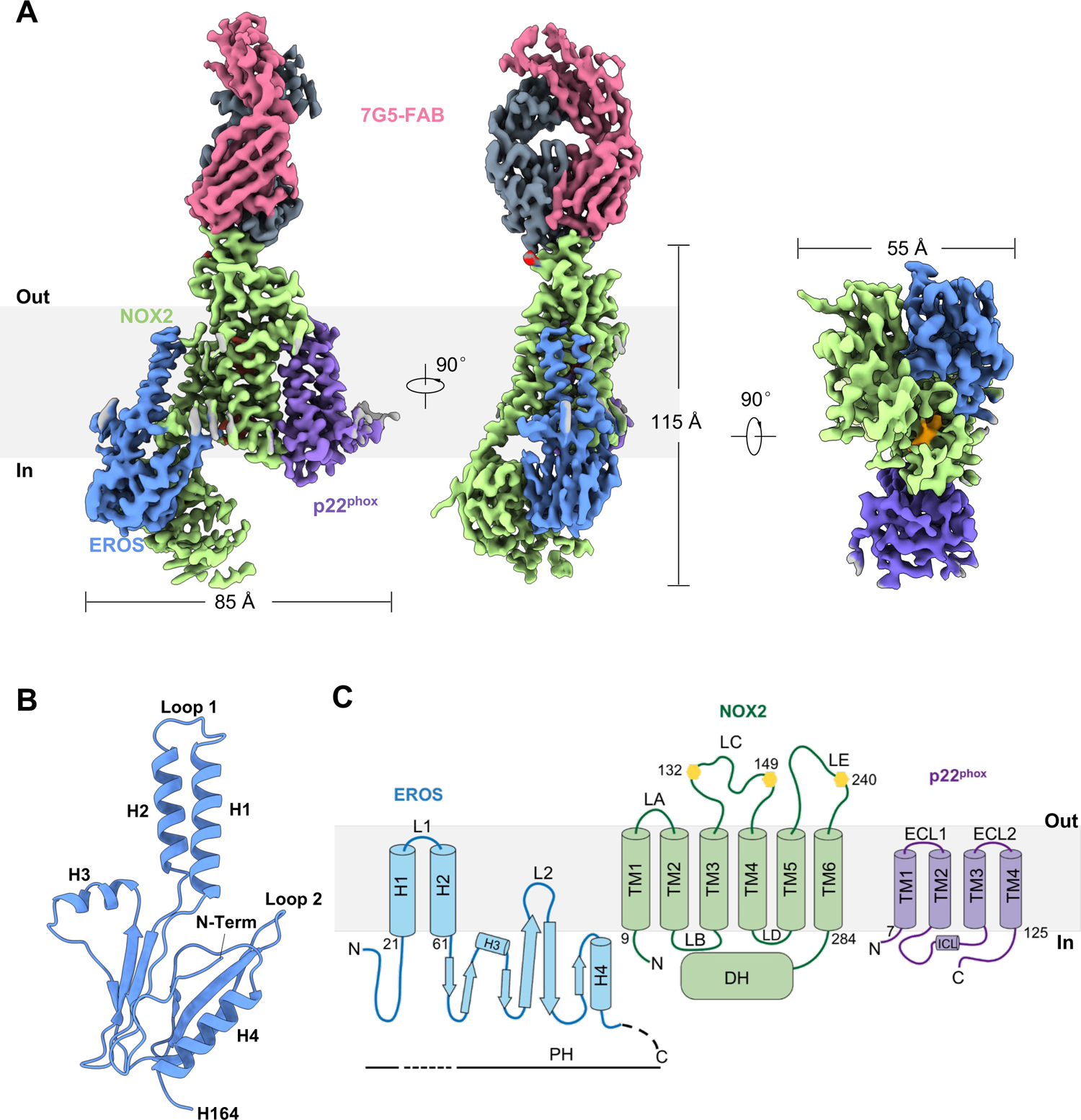
Overall structure of the EROS-NOX2-p22^phox^-7G5 complex. **A.** The cryo-EM map of the EROS-NOX2-p22^phox^-7G5 complex. EROS (blue) and p22^phox^ (purple) are opposite to each other and both associated with NOX2 (green). There is no direct interaction between EROS and p22^phox^. For clarity, the 7G5-Fab dimers are colored differently in pink and gray. The left panel shows a side view of the cryo-EM map, and the middle panel depicts a side view at a 90° rotation to the left panel. The right panel is a bottom view of the complex from an intracellular perspective. The inner heme (orange) is visible in this view. **B.** Cartoon representation of the EROS structure. There are four helices (H1-H4) and six β strands, with the N terminus buried inside and the C terminal fragment (C165-S187) is disordered (not shown). H1 (I21-Y40) and H2 (W47-Q61) are connected by Loop 1; H3 (L85-F93) is nearly perpendicular to H1 and H2. The C terminal H4 (R147-L161) is tilted relative to plasma membrane and does not form a transmembrane helix. The two longest β-strands are anti-parallel and connected by Loop 2 (V117-G121). **C.** Topological model of EROS (blue), NOX2 (green) and p22^phox^ (purple) in plasma membrane. The yellow hexagon labels indicate three glycosylation sites on NOX2. LA-LE corresponds to Loop A to Loop E. DH denotes the dehydrogenase domain of NOX2. PH represents the Pleckstrin Homology domain separated by H1 and H2. L1 and L2 refer to Loop 1 and Loop 2. ECL and ICL stand for extracellular and intracellular loops, respectively.

A molecular model of EROS (Fig. 2B) was built based on the cryo-EM density map. In this model, EROS shows 4 α-helices, of which H1 and H2 separates the Pleckstrin Homology (PH) domain of EROS and is connected by the L1 loop. The two helices (about 4-5 turns) are shorter than the TMs of NOX2 (7-9 turns). The shortest helix (H3) has only 3 turns and is nearly perpendicular to the plane of H1 and H2. The H4 helix is located intracellularly and is only 24 amino acids away from the C-terminal end. Between H2 and H4 there are 6 β-strands, of which the 2 longest β-strands are connected by the L2 loop (Fig. 2C and Fig. S4). These structures constitute the majority of the PH domain that interacts with the cytoplasmic tail of NOX2 as detailed below.

A molecular model of the EROS-NOX2-p22^phox^ complex is shown in Fig. 3A. The two hemes in between TM3 and TM5 of NOX2 are clearly visible. There is also a streak of lipids (light blue) in the NOX2-p22^phox^ interface. There are 3 carbohydrate moieties in the extracellular loops of NOX2 on asparagine residues N132, N149 and N240, consisting with its mature status [18]. H1 of EROS is juxtaposed to TM2 of NOX2, inducing an upward rotational shift of a.a. 67-83 in TM2 for nearly 79° (Fig. 3B) compared with the recently reported NOX2-p22^phox^ structure (PDB ID: 8GZ3). H2 of EROS is closely associated with TM6 of NOX2, inducing a clockwise turn of a.a. 265-292 of TM6 by approximately 48° (Fig. 3C). An extracellular view (Fig. 3D) and intracellular view (Fig. 3E) were provided for better appreciation of these structural changes induced by EROS.

**Figure 3.**
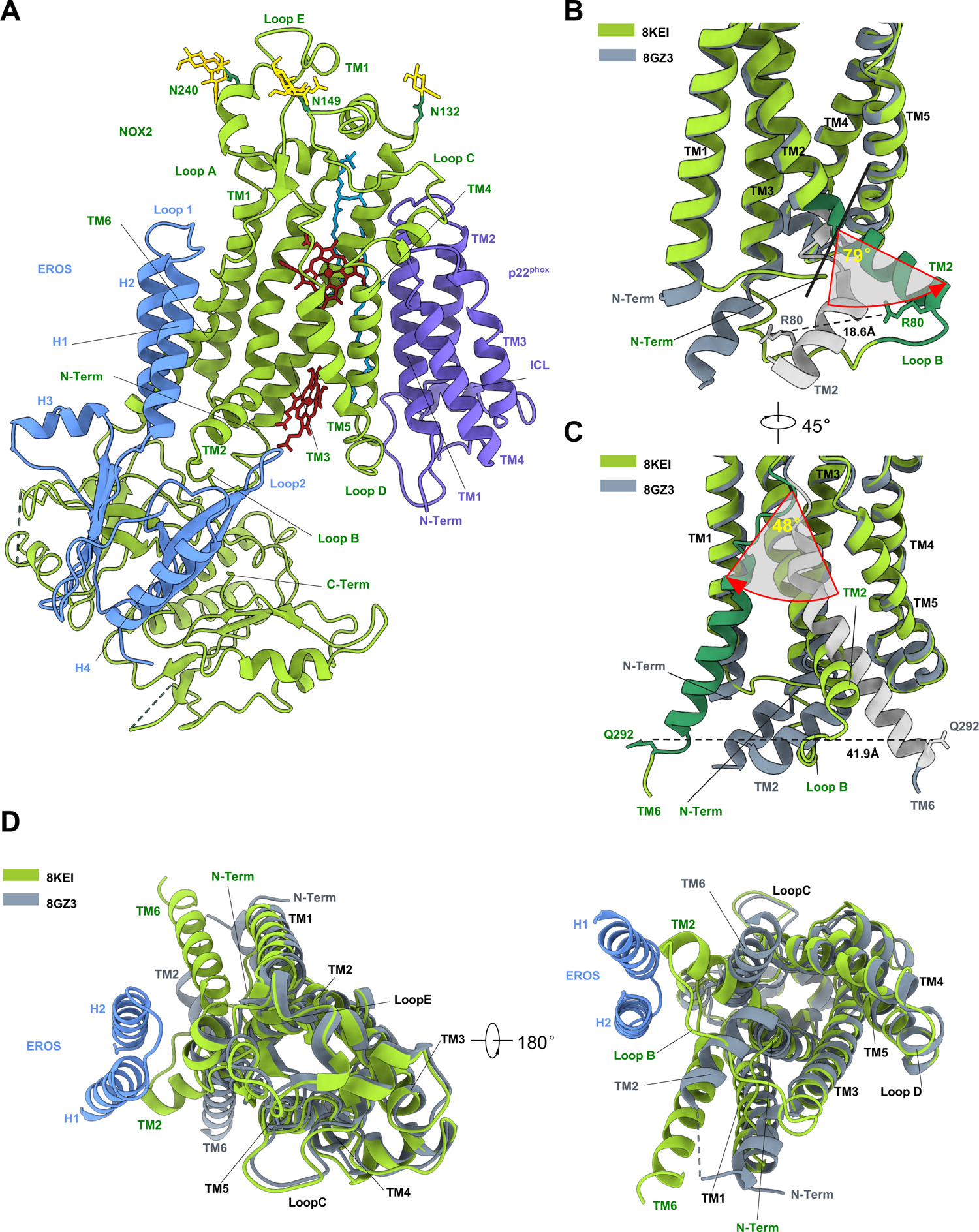
Cartoon representations of the EROS-NOX2-p22^phox^ complex. **A.** The 3-D reconstruction of the EROS-NOX2-p22^phox^ complex, with NOX2 in green, EROS in blue and p22^phox^ in purple. The heme groups are marked in maroon. **B.** TM2 of NOX2 in the presence of EROS (green) vs. in its absence (gray, PDB ID: 8GZ3), showing a nearly 79° upward rotational shift (red arrow) of a.a. 67-83 of TM2 (dark green) from its position in the resting state (silver). The atom-to-atom distance of the shifted R80 is 18.6Å. TM6 and EROS are removed from this panel for clarity. **C.** A 45° horizontal rotation of **B** showing a 48° backward rotation of a.a. 265-292 (dark green) of TM6 when NOX2 is bound to EROS. TM6 in the resting state (PDB ID: 8GZ3) is depicted in gray, and the same fragment in silver. The distance between the shifted Q292 is 41.9Å, along with a movement of the N terminus of NOX2 towards TM2. **D.** Top view (extracellular view, left) and bottom view (intracellular view, right) of the superimposed NOX2 structures in EROS-bound (green) and resting (gray) states, with emphasis on the dislocated TM2 and TM6 of NOX2. EROS is marked in blue.

### The EROS-NOX2 interface

An analysis of the complex structure identified extensive interactions between EROS and NOX2. These include Y51 of EROS that forms a hydrogen bond with TM6 of NOX2, along with the hydrophobic interaction clusters, including A37 on H1, F60/F57 on H2, and S41/P43 on Loop 1, attract and stabilize TM6. R22/L26 on H1 and N62 on H2 interact with TM2 of NOX2, including polar interactions that play an important role in the conformational changes of TM2 (Fig. 4A, 4B). In addition to these interactions that partially stabilize the displaced TM2, there are two clusters of amino acids including L75-R80 on TM2 and G81-C86 on Loop B of NOX2 that interact closely with EROS, including polar interactions between E115/Q140 of EROS and C86/C85 of NOX2 respectively. Together, these interaction clusters form a pocket-like structure surrounding part of TM2 and Loop B of NOX2 (Fig. 4B, 4C), thereby contributing to a marked conformational change in TM2. Moreover, Loop 2 of EROS inserts into a structure that connects Loop B and TM3 of NOX2, forming a hydrogen bond between Y119 of EROS and the inner heme (Fig. 4B, 4C). There is also an interaction between R118/Y119 of EROS and W206/H210 on TM5 of NOX2 near the inner heme. As a result of these interactions, the position of the inner heme was shifted downward by a few angstroms, and the metal-to-metal distance between the two hemes were extending to 21.1Å from 19.8Å in the absence of EROS [17].

**Figure 4.**
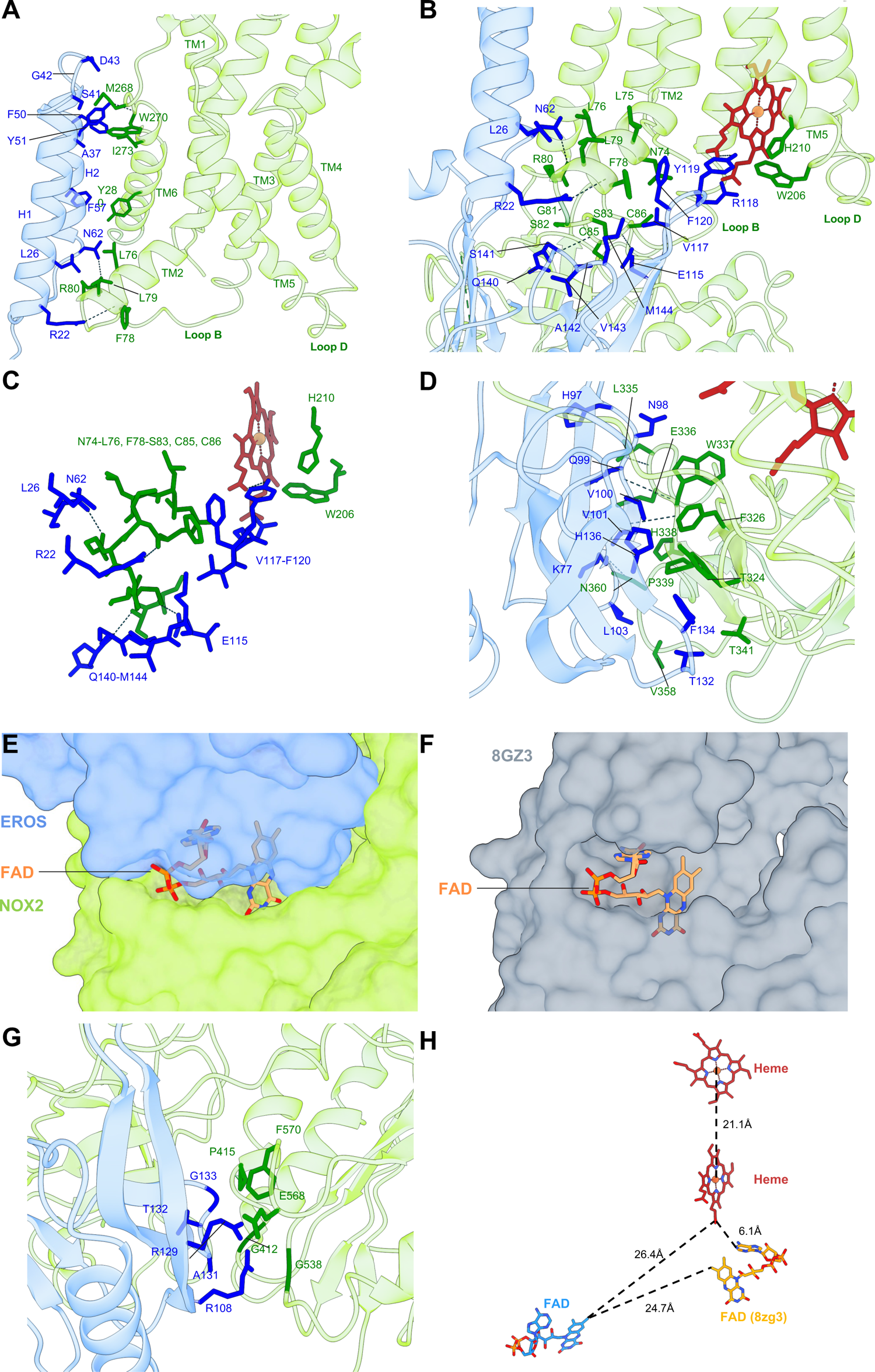
The EROS-NOX2 interface. **A.** Interactions between H1 and H2 of EROS and TM2 and TM6 of NOX2. Of note, EROS residues R22 and N62 form hydrogen bonds with NOX2 residues F78 and R80, respectively. Additionally, the EROS residue Y51 is involved in hydrogen bonding with NOX2 residue W270. Residues at this interface (EROS: R22, L26, A37, S41, p43, F50, Y51, F57, N62; NOX2: L76, F78, L79, R80, M268, W270, I273, Y280). The residues of EROS and NOX2 are depicted as blue and green sticks, respectively; the hydrogen bonds are represented by dark dash lines. **B.** Interaction of the inner heme with EROS. Loop 2 protrudes from EROS and forms a hydrogen bond between Y119 and the lower heme (maroon). Two other EROS residues, E115 and Q140, form hydrogen bonds with NOX2 residues C86 and C85 on Loop B, respectively. In addition, there are hydrophobic interactions between R118/Y119 of EROS and W206/H210 of NOX2 in TM5. **C.** The TM domains are removed to show more clearly pocket-like interactions between NOX2 and three clusters of EROS including four polar interactions (R22, N62, E115, Q140 on EROS). **D.** The EROS-NOX2 interface in the FAD-binding domain of NOX2. A total of 20 amino acids, 10 each from EROS and NOX2, participate in this interaction and form a tight zipper that prevents FAD access to the DH domain of NOX2. The hydrogen bonds are shown as dark dashed lines. **E and F.** Cartoon and surface representation of FAD binding to NOX2 (green) in the presence of EROS (blue in **E**) and in its absence (**F**, NOX2 is shown in gray). **G.** The EROS-NOX2 interface in the NADPH-binding domain of NOX2. Ten residues, five from EROS (R108, R129, A131, T132, G133) and the other five from NOX2 (G412, P415, G538, E568, F570) are involved in this interaction. **H.** Schematic representation of the electron transfer path showing the relative positions and distances between FAD and the two hemes in the absence (FAD in orange) and presence (FAD in blue) of EROS. The edge-to-edge distance between the inner heme and FAD in resting NOX2 (6.1 Å) is shorter than that in EROS-NOX2 (26.4 Å). The ferric ions are shown as orange spheres.

The cytoplasmic dehydrogenase homology (DH) domain houses two structural subdomains for binding of FAD and NADPH, respectively [6, 19]. The importance of the DH domain in the initiation of transmembrane electron transfer is evidenced by 19 missense mutations in this region that cause CGD [19]. An analysis of the EROS-NOX2 interface identified close interactions between the PH domain of EROS and the DH domain of NOX2 (Fig. 4D-4G). These interactions can be roughly divided into two categories. The first involves a subdomain where NOX2 binds to FAD and includes T324, L335, E336, W337, H338, P339, T341, V358, and N360 of NOX2. In this subdomain, L335/W337 could form hydrogen bond with Q99 of EROS; there is also a polar interaction between W337 and V101 of EROS. Three amino acids in this subdomain, H338, P339, and T341 have been reported to be substituted due to missense mutations in CGD patients [19], indicating that these amino acids play a key role in FAD binding. The presence of EROS may interfere with FAD binding to the DH domain, as FAD was not observed in our cryo-EM structure. Consistent with this prediction, when we fitted our model with that of the FAD-bound NOX2 (PDB ID: 8ZG3), three EROS amino acids (V101, L103, and F134) conflicted with FAD binding (Fig. 4D). These findings provide strong evidence that EROS interaction with NOX2 prevents FAD access to the DH domain (Fig. 4E, 4F).

The second region in the DH domain is responsible for NOX2 binding to its substrate NADPH. In this region, the amino acids G412/P415/G538/E568/F570 of NOX2 form tight hydrophobic interactions with R108/R129/A131/T132/G133 of EROS (Fig. 4G). Previous studies have shown that the affinity of resting NOX2 for NADPH is lower than that for FAD [20]; in the presence of EROS, the ability of NOX2 to bind NADPH could be further compromised based on our structural analysis. Moreover, missense mutations of G412, P415, and E568 have been reported in CGD patients [5], indicating the importance of this subdomain of NOX2 in binding NADPH.

The conformation of NOX2 was markedly altered in the presence of EROS, resulting in an increased distance between the two hemes (21.1Å) compared with a previously reported structure (PDB ID: 7U8G, 19.8Å). The original FAD binding subdomain moved away from the inner heme site (compared to 8ZG3). Structurally, EROS occupies the FAD and NAPDH binding subdomains in NOX2. Spatially, there is a shift of the FAD binding subdomain away from where it used to be (Fig. 4H). These structural changes suggest that EROS binding interferes with the electron transport chain of NOX2 and compromises superoxide production by NOX2.

### The EROS-bound NOX2 is in an inactive state

Analysis of the structure of the EROS-NOX2-p22^phox^ suggests that EROS association with NOX2 prevents FAD access to the DH domain, hence altering the electron transfer pathway. If this is the case, the inhibitory mechanism must be alleviated for normal transfer of electron through the active NOX2. To determine whether EROS association is dynamic with respect to NOX2 activation, a NanoLuc complementation assay [14, 21] was adopted to measure the dynamic changes in distance between the C terminal end of EROS (fused with LgBiT) and N terminal end of NOX2 (fused with SmBiT). The expression plasmids containing these engineered EROS and NOX2 fusion constructs (Fig. 5A) were cotransfected into COS-7 cells together with plasmids expressing the necessary cytosolic factors for reconstitution of active NOX2 (Fig. 5B). Upon stimulation with phorbol myristate acetate (PMA), that induces phagocyte superoxide production, the changes in luminesce were measured in real time that reflect the degree of association between EROS and NOX2. As shown in Fig. 5C, PMA induced a time-dependent dissociation of EROS from NOX2. Under the same stimulation conditions, PMA induced superoxide production in the reconstituted COS-7 cells (COS^91/22^) that were significantly inhibited by co-expression of EROS (Fig. 5D). Together, these results suggest that EROS binding keeps NOX2 in an inactive state.

**Figure 5.**
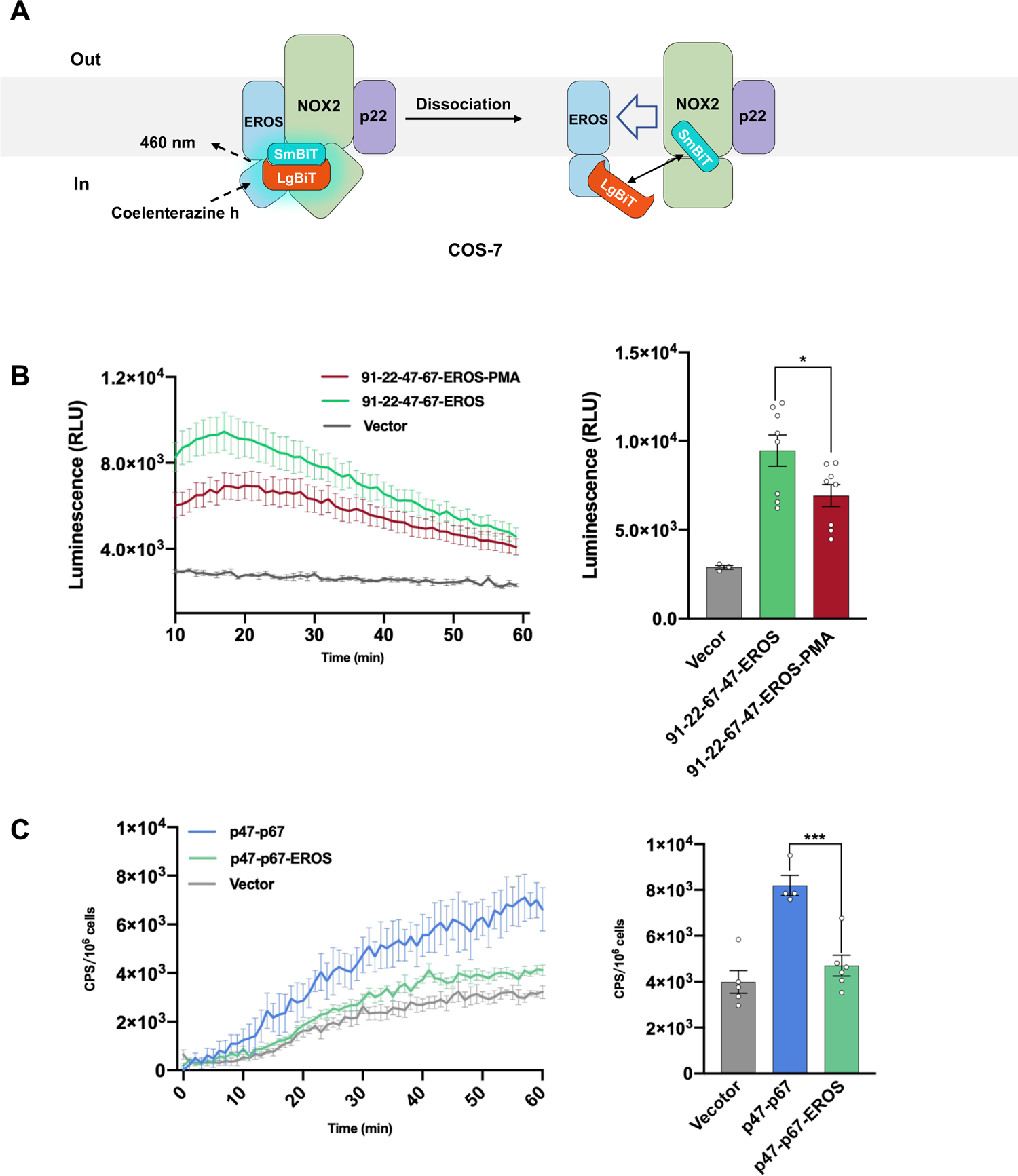
The EROS-bound NOX2 in an inactive state. **A.** Schematic representation of NanoLuc complementation assay. The two components, LgBiT and SmBiT, were fused to EROS and NOX2 respectively, as detailed in *Methods*. These engineered constructs (EROS-N-LgBiT, NOX2-C-SmBiT) were cotransfected into COS-7 cells together with the p47^phox^ and p67^phox^ expression plasmids. The changes in luminescence intensity were measured after addition of the substrate coelenterazine H (10 μM) and PMA (200 ng/ml), with a 1 min interval recording at 460 nm. Hypothetical dissociation of EROS from NOX2 is marked by an arrowhead. **B.** Data from 10 to 60 min after PMA addition are shown and quantified in the right panel. **C.** Superoxide production in reconstituted COS-7 cells (COS^91/22^) cotransfected with expression plasmids of p47^phox^, p67^phox^, with or without an EROS expressing plasmid. The PMA-induced superoxide production was measured in real time for 60 min using isoluminol, and data were quantified and presented in the right panel. Data shown are mean ± SEM based on multiple independent experiments. *, *p* < 0.05, ***, *p* <0.001.

## DISCUSSION

In this study, we present the first high-resolution structure of EROS bound to NOX2-p22^phox^, based on cryo-EM electron density maps. The structure covers the entire EROS protein except the C terminal 23 amino acids that appears to be disordered and not involved in binding NOX2. There is no direct interaction between EROS and p22^phox^ because these two proteins are placed on opposite sides of NOX2. The first 2 α-helices of EROS are anti-parallel and membrane embedded with ∼5.5 turns (H1) and ∼4.5 turns (H2), respectively. Although shorter than the TMs in NOX2, these α-helices appear to form transmembrane domains as the molecule can be detected in intact cells by an anti-EROS antibody. EROS associates with the TM helices of NOX2 through H1 (interacting with TM2) and H2 (interacting with TM6), causing conformational changes of both TM helices. For the remaining structure of EROS, helices H3 and H4 do not directly interact with NOX2, but the cytoplasmic β-strands form extensive interactions with the cytoplasmic domains of NOX2 including the DH domain that harbors the FAD and NADPH binding sites. These interactions are expected to block access of FAD and NADPH to NOX2. The solved cryo-EM structure of the EROS-NOX2-p22^phox^ complex does not contain FAD or NADPH. There is also a pocket-like structure on EROS that interacts with part of the TM2 and Loop B of NOX2. This loop between TM2 and TM3 is also very close to the L2 loop of EROS that forms a hydrogen bond with the inner heme. Altogether, these interactions alter the electron transfer pathway by increasing the distance between the heme groups, inducing a shift of the FAD binding site, and blocking access of FAD and NADPH to the DH domain of NOX2.

The discovery of EROS as a molecular chaperone of NOX2 [11] is a major advancement in NOX2 biology and adds a new etiological factor for CGD [12, 13, 22]. Through a series of elegant experiments, Thomas and coworkers showed that EROS associates in the endoplasmic reticulum (ER) with the immature (58 kDa) NOX2 polypeptide and prevents its degradation [14]. Although EROS is considered an ER resident protein, its primary sequence does not contain a typical ER-retention motif [23, 24]. EROS associates with NOX2 most probably before it incorporates the heme groups [14], whereas p22^phox^ binds NOX2 after heme incorporation and N-glycosylation with high-mannose [25, 26]. These observations indicate that NOX2 polypeptide first forms a heterodimer with EROS and then the complex becomes a heterotrimer with the addition of p22^phox^. Both EROS and p22^phox^ play important roles for NOX2 maturation from precursor (65 kDa) for efficient expression to plasma membrane [14, 25]. Our finding that EROS associates with mature NOX2 bearing complex oligosaccharides (91 kDa) suggests that the EROS-NOX2-p22^phox^ complex has gone through the Golgi apparatus where NOX2 is fully glycosylated [26]. Indeed, our structural model of NOX2 shows carbohydrate moieties at the expected N132, N149 and N240, respectively [18]. Of interest, previous work involving the purification of NOX2-p22^phox^ heterodimer (flavocytochrome *_b558_*) did not identify the presence of EROS, and there might be several reasons. First, the detergents used for purification of neutrophil flavocytochrome b558, such as Triton X-100 and octylglucoside [27], are quite strong for separation of EROS from flavocytochrome b558. In our study, mild detergents including LMNG and CHS were used in order to obtain the EROS-NOX2-p22^phox^ heterotrimeric complex for cryo-EM analysis. Secondly, the size of EROS (187 a.a. predicted m.w. of 20,774 Da) is very close to that of p22^phox^ (195 a.a. predicted m.w. of 21,013 Da) and they appear in SDS-PAGE as partially overlapping bands. Thirdly, there might be signs for the presence of EROS that were not further pursued at that time. For example, chemical cross-linking of purified components of flavocytochrome b558 resulted in a complex with m.w. 120-135 kDa on SDS [27], which is larger than the size of NOX2 (91 kDa) and p22^phox^ (22 kDa) combined. Given the 1:1 stoichiometry of NOX2 and p22^phox^ in the heterodimer [28], there might be another component of 20-22 kDa in the cross-linked complex. Finally, EROS is widely distributed but does not express in large quantity, and it only appeared in complex with NOX2-p22^phox^ in our experiments. These factors may have contributed to the delays in finding EROS as a NOX2-binding protein.

NOX2 is the catalytic subunit of phagocyte NADPH oxidase that contains all the necessary components for one-electron reduction of molecular oxygen, including two nonidentical hemes, binding sites for the cofactor FAD and the substrate NADPH [6, 7, 20, 26]. In comparison, current models do not support any catalytic activity of p22^phox^ and EROS in the heterotrimeric complex [6]. In addition to providing a docking site for p47^phox^ membrane translocation [29–31], p22^phox^ has been known to stabilize the newly synthesize and heme-incorporated NOX2 through heterodimeric complex formation.

With the discovery of EROS as a molecular chaperone of NOX2 that forms a heterodimer with NOX2 prior to p22^phox^, it appears that EROS and p22^phox^ work together to stabilize the newly synthesized NOX2 precursor and ensure its maturation and proper transit to plasma membrane. Both EROS and p22^phox^ associate with NOX2 with their TM helices. There are, however, significant differences between EROS and p22^phox^ in their interaction with the C terminal cytoplasmic fragment of NOX2. Whereas recently published NOX2 structural models [16, 17] and the present study do not show any interaction between the cytoplasmic domains of NOX2 and p22^phox^, our work provides direct evidence for close interaction between EROS and NOX2 in their C terminal cytoplasmic fragments including formation of multiple hydrogen bonds. These interactions may be necessary for stabilization of NOX2 as it matures from ER to Golgi, but it is also important to note that the C terminal fragment of NOX2 contains binding sites for FAD and NADPH. Previously studies have identified several missense mutations of amino acids in this region that cause CGD through disruption of the electron transfer pathway [19]. It is therefore speculated, based on structural information provide in this study, that the EROS-bound NOX2 remains inactive for not being able to bind FAD and NADPH (Fig. 6A). This ‘inactive state’ is conceptually different the from previously proposed ‘resting state’, which emphasizes on activation by the translocated cytosolic factor and Rac-GTP. Instead, our proposed working model focuses on the release of inhibition by EROS, thus emancipating the catalytic capability of NOX2 (Fig. 6B, 6C). It was previously shown that the purified and relipidated flavocytochrome b558 was able to generate superoxide in a FAD-and NADPH-dependent, but cytosolic factor-independent manner [32]. Given the high affinity of FAD binding to the DH domain and the abundance of NADPH in the cytosolic compartment, regulation of superoxide production must include a preventive measure. Our NanoLuc complementation results have shown that PMA, a stimulant for superoxide production, can induce dissociation of EROS from NOX2 in reconstituted cells. It is therefore possible that the signals for EROS dissociation come from the same stimulants that activate the NADPH oxidase. In this regard, dissociation of EROS from NOX2 may be coupled to the assembly of the active NOX2 complex (Fig. 6).

**Figure 6.**
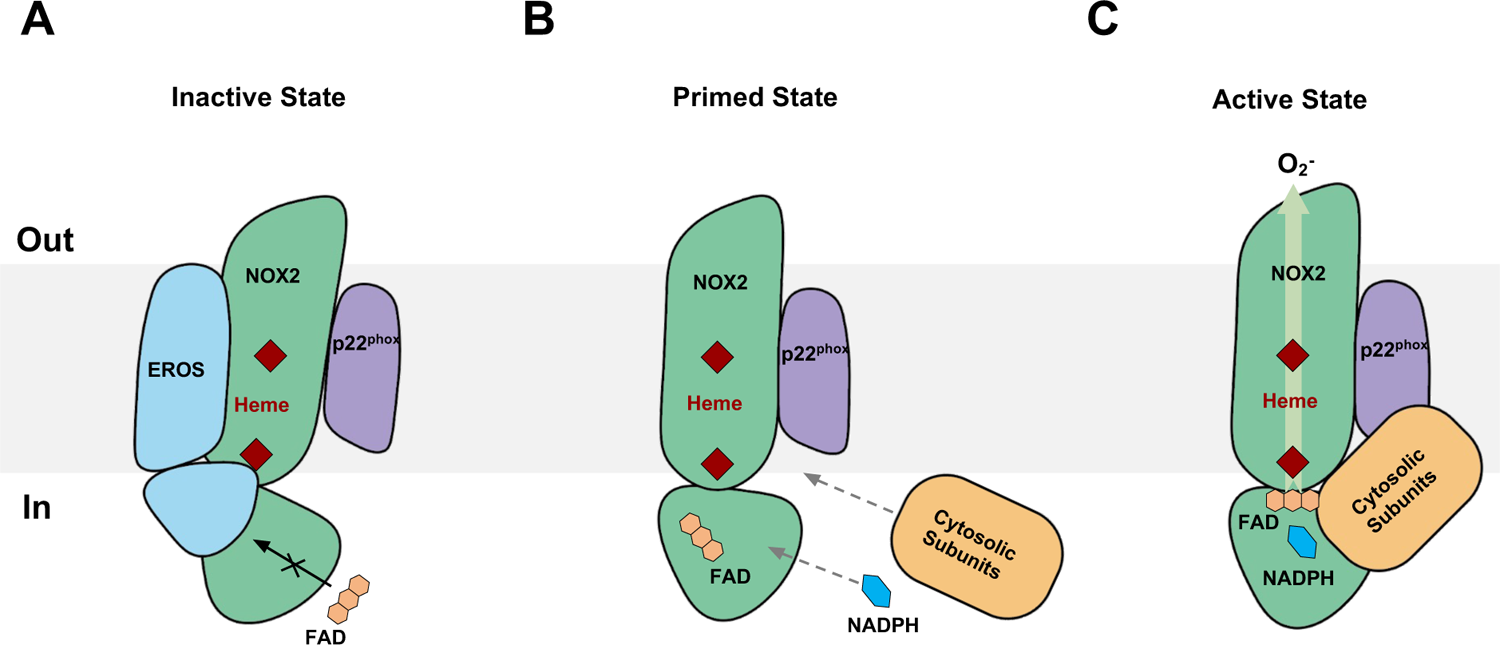
A hypothetical model of NOX2 activation taking EROS into consideration. **A.** NOX2 in the inactive state. The association of EROS protects nascent NOX2 against proteosomal degradation and, at the same time, prevents FAD and NADPH from binding to the DH domain. **B.** NOX2 in the primed state. Priming signals induce dissociation of EROS from NOX2, allowing FAD access to NOX2. **C.** NOX2 in the active state. Docking of the cytosolic NOX activators and Rac-GTP changes the conformation of NOX2, permitting NADPH binding and electron transfer.

In summary, the solved structure of EROS-NOX2-p22^phox^ provides previously unavailable information on the interaction between EROS and NOX2. EROS is closely associated with NOX2 not only through the transmembrane helices but also through the intracellular domains, potentially preventing FAD from binding to the DH domain of NOX2. Our work complements the structural analysis of the NOX2-p22^phox^ heterodimeric complex in the absence of EROS [16, 17], what apparently shows an intact electron transfer pathway. In addition, the present study provides a structural basis for EROS binding to NOX2, that was reported in several recent publications [11, 14] and was associated with newly identified cases of CGD [11–13, 22]. Based on our structural model, we propose that EROS holds NOX2 in an inactive state so as to prevent inadvertent production of superoxide. Additional work will be required to confirm the inhibition of NOX2 activity by EROS and delineate the mechanisms for the dissociation of EROS from the DH domain of NOX2 and initiation of electron transfer through the catalytic subunit.

## METHODS

### Plasmid vector construction

Wildtype human NOX2 and N-terminal FLAG tag-p22^phox^ were cloned into the pMlink vector [33] to generate pMlink-N-FLAG-p22^phox^-NOX2. The DNA coding sequences for wildtype human EROS, 7G5 Fab heavy chain and light chain were synthesized by Genewiz (Suzhou, China). These cDNAs were then cloned into other pMlink vectors for co-expression of the relevant proteins. The plasmid construct of NOX2 used in fluorescent confocal microscopy was prepared by fusion of the green fluorescent protein Clover coding sequence to the cDNA of NOX2 without the initiation codon, and the fused cDNA was cloned into the pcDNA3.1 expression plasmid (Invitrogen, Calsbad, CA). For visiualization of EROS, the coding sequence of the red fluorescent protein mRuby2 was fused to the C terminal end of EROS and the resulting cDNA was cloned into pcDNA3.1. For NanoBiT-based assay, the LgBiT coding sequence was fused to the N terminus of EROS to generate pcDNA3.1-N-LgBiT-EROS. The SmBiT coding sequence was fused to the C terminus of NOX2 coding sequence to generate pCDNA3.1-NOX-C-SmBit. The cDNAs of human p47^phox^ and p67^phox^ were cloned into pcDNA 3.1. All constructs were verified by DNA sequencing using the service of Genewiz.

### 7G5 Fab purification

7G5 Fab [16] were expressed in transiently transfected Expi-293F cells (Thermo Fisher Scientific, Waltham, MA) and purified by affinity chromatography using protein A/G agrose (Beyotime, Shanghai, China). The elution was collected and incubated with EROS-NOX2-p22^phox^ preparation for complex formation.

### Complex purification

Expi-293F cells were cultured in FreeStyle 293 medium (Union Biotech, Shanghai, China) at a density of 2.0×10^6^ cells/ml and co-transfected with pMlink-N-FLAG-p22^phox^-NOX2 and pMlink-EROS using PEI 4000 reagent (Polysciences, Warrington, PA). The transfected cells were cultured for 48 h and collected by centrifuged (4,000 × g, 15 min). The cell pellet was washed with Buffer A (20 mM Tris pH 8.0, 150 mM NaCl, 1 mM phenylmethanesulfonyl fluoride (PMSF), 1 μg/ml L/P/A (leupeptin/pepstatin/aprotinin)), and resuspended in Buffer B (20 mM Tris pH 8.0, 150 mM NaCl, 20% (v/v) glycerol, 1% Lauryl maltose neopentyl glycol (LMNG), 0.1% cholesteryl hemisuccinate (CHS), 1mM PMSF, 1 μg/ml L/P/A) and lysed for 2 h at 4°C to solubilize membrane proteins. The resulting supernatant containing soluble membrane proteins was collected by centrifugation at 18,000 rpm for 60 min. The supernatant was then incubated with 2-5 ml of FLAG Affinity resin (GenScript, Nanjing, China) for 1 h at 4°C. Subsequently, the resin was washed with Buffer C (20 mM Tris pH 8.0, 150 mM NaCl, 0.01% LMNG and 0.001% CHS) and incubated with elution buffer (20 mM Tris pH 8.0, 150 mM NaCl, 0.01% LMNG, 0.001% CHS, 200 μg/ml 3 X FLAG peptide) for 30 min. The elution was collected and subsequently incubated with 7G5 Fab for another 30 min. The EROS-NOX2-p22^phox^-7G5 Fab complex was concentrated and applied onto a Superose 6 Increase 10/300 GL column (GE HealthCare, Chicago, IL) in Buffer D (20 mM Tris pH 8.0, 250 mM NaCl, 0.06% digitonin) for size-exclusion chromatography (SEC). The fractions with the highest protein concentration were used for cryo-EM sample preparation and SDS-PAGE.

### Cryo-EM grid preparation and data collection

For cryo-EM grid preparation, 3 μL of the purified complex (8 mg/ml) was deposited onto a glow-discharged holey grid (Quantifoil Au 300 mesh R1.2/1.3, Quantifoil, Großlöbichau, Germany). It was then plunge-frozen in liquid ethane using the Vitrobot Mark IV (Thermo Fischer Scientific). Cryo-EM images were obtained at the Kobilka Cryo-EM Center of The Chinese University of Hong Kong, Shenzhen, on a Titan Krios C3i cryo-TEM (Thermo Fisher Scientific) operating at 300 kV, using a K3 Summit detector (Gatan, Pleasanton, CA) with a pixel size of 0.85 Å. Inelastically scattered electrons were eliminated by a GIF Quantum energy filter (Gatan) using a slit width of 20 eV. Images were captured at defocus values ranging from −1.2 to −2.5 μm using the semi-automatic data acquisition software, SerialEM (University of Colorado, Boulder, CO). A total of 12,322 image stacks were collected within 96 h, with a total dose of 52 electrons per square angstrom over 2.5 seconds exposure on each movie.

### Cryo-EM image processing and map construction

The cryoSPARC v3.3.1 software (Structura Biotechnology, Toronto, Canada) was utilized to perform single particle analysis of the EROS-NOX2-p22^phox^-7G5 complex. The image stacks underwent motion correction and patch CTF estimation. A total of 3,470,723 particles were auto-picked and subsequently underwent 2D classification. Based on these results, 1,985,600 particles were selected for *ab initio* reconstruction. After multiple rounds of refinement, a final set of 131,926 particles underwent non-uniform refinement and local refinement, resulting in a map with an overall resolution of 3.56 Å at a Fourier shell correlation of 0.143.

### Model building and refinement

The initial model for rebuilding and refinement against the electron microscopy density map was the model of the resting-state NOX2 (PDB: 8GZ3). The electron microscopy density map was docked with the model using UCSF Chimera-1.14, followed by iterative manual adjustment and rebuilding in COOT-0.9.8. The models were further refined and validated using the Phenix-1.20 programs (see Supplementary Table 1). Structural figures were generated using UCSF Chimera-1.14, ChimeraX-1.27, and PyMOL-2.5.

### Flow cytometry analysis

HL-60 cells differentiated with 1.3% DMSO for 6 days (dHL-60) were collected by centrifugation at 500 x *g* for 5 min, the cells were harvested and washed with 1X HBSS. Subsequently, the cells were blocked on ice with 5% BSA in HBSS for 30 min.

Fluorescent antibody staining was then conducted on ice. The cells were stained with an anti-EROS FITC-conjugated antibody (CSB-PA859832LC01HU; CUSABIO Technology, Houston, TX, used at 1:20 dilution with HBSS) or an anti-gp91^phox^ 7D5 antibody (D162-3; Medical & Biological Laboratories, Nagoya, Japan, used at 1:200 dilution in HBSS) with a fluorescent secondary antibody, or both anti-EROS FITC and anti-gp91^phox^ 7D5. The goat anti-mouse PE secondary antibody (12-4010-82; Thermo Fisher Scientific) was used at a dilution of 1:100 in HBSS. After incubation on ice for 30 min, the cells were washed and suspended in HBSS, and analyzed using flow cytometry on a CytoFLEX S flow cytometer (Beckman Coulter, Brea, CA). The FlowJo™ v10.8 Software (BD Life Sciences) was used for flow cytometry data analysis. Single-staining samples were used to configure gating and compensation for FITC and PE channels.

### Fluorescent confocal microscopy

For confocal microscopy, dHL-60 cells were centrifuged at 800 x g for 5 min, and the collected cells were washed 2-3 times with PBS. An appropriate amount of cell suspension was pipetted onto a slide and then fixed with 4% paraformaldehyde for 20 min. The cells were rinsed with PBS three times, and incubated with anti-EROS-FITC and anti-NOX2-PE antibodies overnight at 4°C. The cells were again rinsed with PBS three times and then incubated with DAPI for 5 min. Finally, a sealer containing an anti-fluorescence quencher was added to seal the slide.

COS-7 cells were cultured for 12 h on glass coverslips in a 24-well plate and then co-transfected with pcDNA3.1-NOX2-N-Clover and EROS-C-mRuby2 using Lipofectamine 3000 (Thermo Fisher Scientific) for 24 h at 37°C. The cells were rinsed with PBS and fixed with 4% paraformaldehyde for 10 min at room temperature. The coverslips were then mounted on glass slides and imaged using a confocal microscope (LEICA TCS SP8, Leica Microsystems, Buffalo Grove, IL) with a 40 X oil objective.

### Superoxide production assay

COS^91/22^ was prepared using the original protocol of Dinauer and coworkers [34] except that the p22^phox^ cDNA was expressed in pcDNA3.1-neuromycin vector (Thermo Fisher Scientific). Superoxide production in COS^91/22^ cells was assessed using an isoluminol-enhanced chemiluminescence (ECL) assay in 6 mm wells of 96-well, flat-bottomed white tissue culture plates. Enzyme-free cell-dissociation buffer (Thermo Fisher Scientific) was used to harvest COS^91/22^ cells that were subsequently washed once using 0.5% BSA/HBSS. Cells were resuspended in 0.5% BSA/HBSS at a density of 1-3×10^6^ cells/ml. Following 5 min of preincubation at room temperature in the dark with 100 μM isoluminol and 40 U/ml HRP, a 200 μL cell aliquot was added to the plate well. The cells were stimulated with PMA (200 ng/ml). A chemiluminescence assay was conducted for continuous measurement of chemiluminescence counts per second (cps) at 37°C in an Envision multimode plate reader (PerkinElmer Life Sciences, Waltham, MA), recording measurements at 1 min intervals.

### NanoBiT based assay

COS-7 cells were seeded in a 6-well plate and co-transfected with pcDNA3.1-N-LgBiT-EROS, pcDNA3.1-NOX2-C-SmBit, pcDNA3.1-p47^phox,^ and pcDNA3.1-p67^phox^ or empty vector using Lipofectamine 3000 for 24 h at 37°C. The cells were washed with HBSS, harvested and then plated onto white 96-well plates at a concentration of 2×10^5^ cells per well. The transfected cells were stimulated with either PMA (200 ng/ml) or buffer control. Next, coelenterazine H (Yeasen Biotechnology, Shanghai, China) is added to the plates at a final concentration of 5 μM. After a 10-minute incubation at 37°C, luminescence was measured on the plate using an Envision multimode plate reader for 50 min.

### Statistical analysis

The Prism software (ver. 8.0, GraphPad, San Diego, CA) was used for statistical analysis of data. For statistical comparison, one-way ANOVA was used. A *p* value of < 0.05 was considered statistically significant.

## AUTHOR CONTRIBUTIONS

S. L. and R.D.Y. designed research and initiated project; S. L. and Y. Z. purified proteins and prepared samples for cryo-EM analysis; F. W. and Z. L. collected cryo-EM data; A. L. built and refined the structural model; S. L. and Y. L. performed functional assays and validated the model; R.D.Y, R.R., Z.L. and T. W. supervised research; S. L., Y. L. and R.D.Y. analyzed data and wrote manuscript with input from other co-authors.

## ACKNOWLEDGEMENTS

The authors thank the High-Performance Computing Platform at The Chinese University of Hong Kong, Shenzhen for computational work. This study was supported in part by grants from the National Key R&D Program of China (Grant 2019YFA0906003); the Science, Technology and Innovation Commission of Shenzhen Municipality Grant GXWD20201231105722002-20200831175432002 (to R.D.Y.); National Natural Science Foundation of China Grant 32070950 (to R.D.Y.); China Postdoctoral Science Foundation 2022M713049 (to A. L.) and Shenzhen Science and Technology Program Grant No. RCBS20221008093330067 (to A. L.); The Kobilka Institute of Innovative Drug Discovery at The Chinese University of Hong Kong, Shenzhen (Z.L., R.R., R.D.Y.); and the Ganghong Young Scholar Development Fund (S.L., Y.Z., R.D.Y.).

---

**Supplemental Figure S1.**
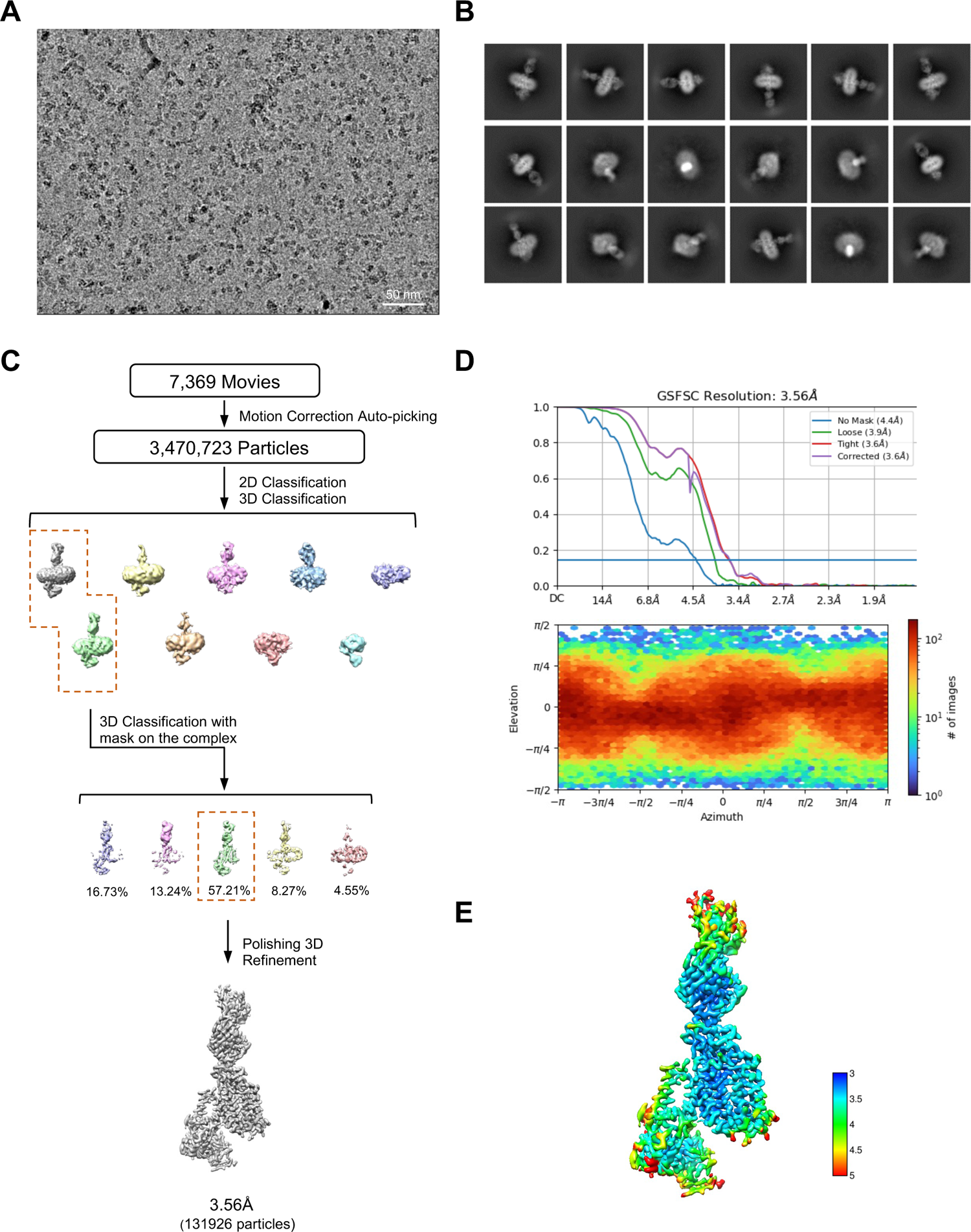
Cryo-EM data processing of the NOX2-EROS complex. **A.** Representative micrograph after motion correction and dose weighting. **B.** 2D class averages of the NOX2-p22^phox^ complex bound with EROS. **C.** Workflow of cryo-EM data processing using cryoSPARC. **D.** Gold standard Fourier shell correlation (FSC) curve indicates an overall nominal resolution at 3.56 Å. **E.** Local resolution map.

**Supplemental Figure S2.**
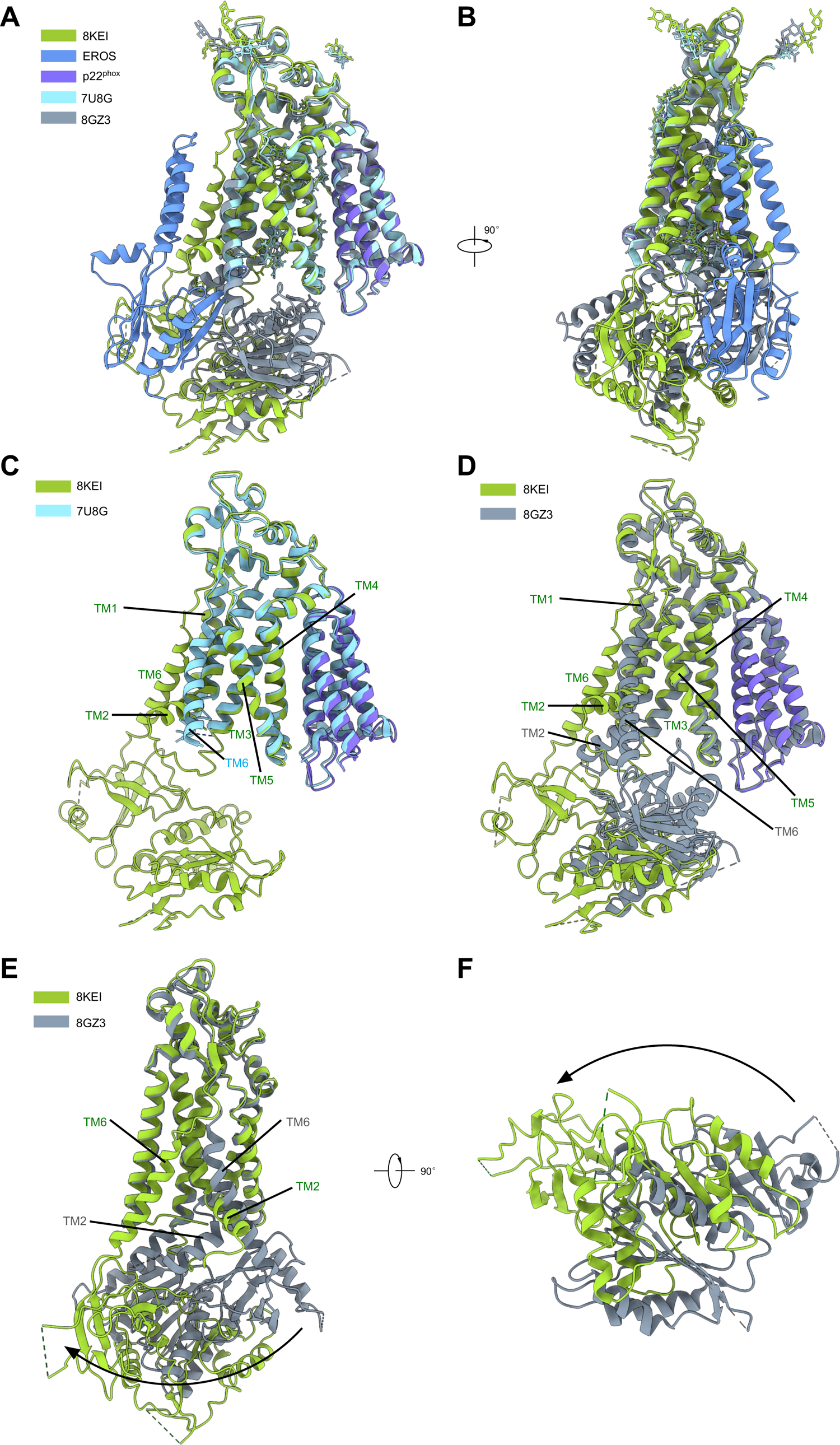
Comparison of EROS-NOX2-p22^phox^ structure to human NOX2-^p22phox^ complex. **A.** Side view of the structural alignments of NOX2 in the inactive state (NOX2: yellow green, EROS: blue, p22^phox^: purple), 8GZ3 (dark gray), and 7U8G (light blue) in cartoon representation. **B.** A 90°-rotated view of **A** is illustrated. **C.** Side view of the structural alignment of NOX2 in the inactive state (NOX2: yellow green, EROS: blue, p22^phox^: purple) and 7U8G (light blue). **D.** Side view of the structural alignment of NOX2 in the inactive state (NOX2: yellow green, EROS: blue, p22^phox^: purple) and 8GZ3 (dark gray). **E.** The movements of DH domain of NOX2 compared to 8GZ3. **F.** A 90°-rotated view of **E**.

**Supplemental Figure S3.**
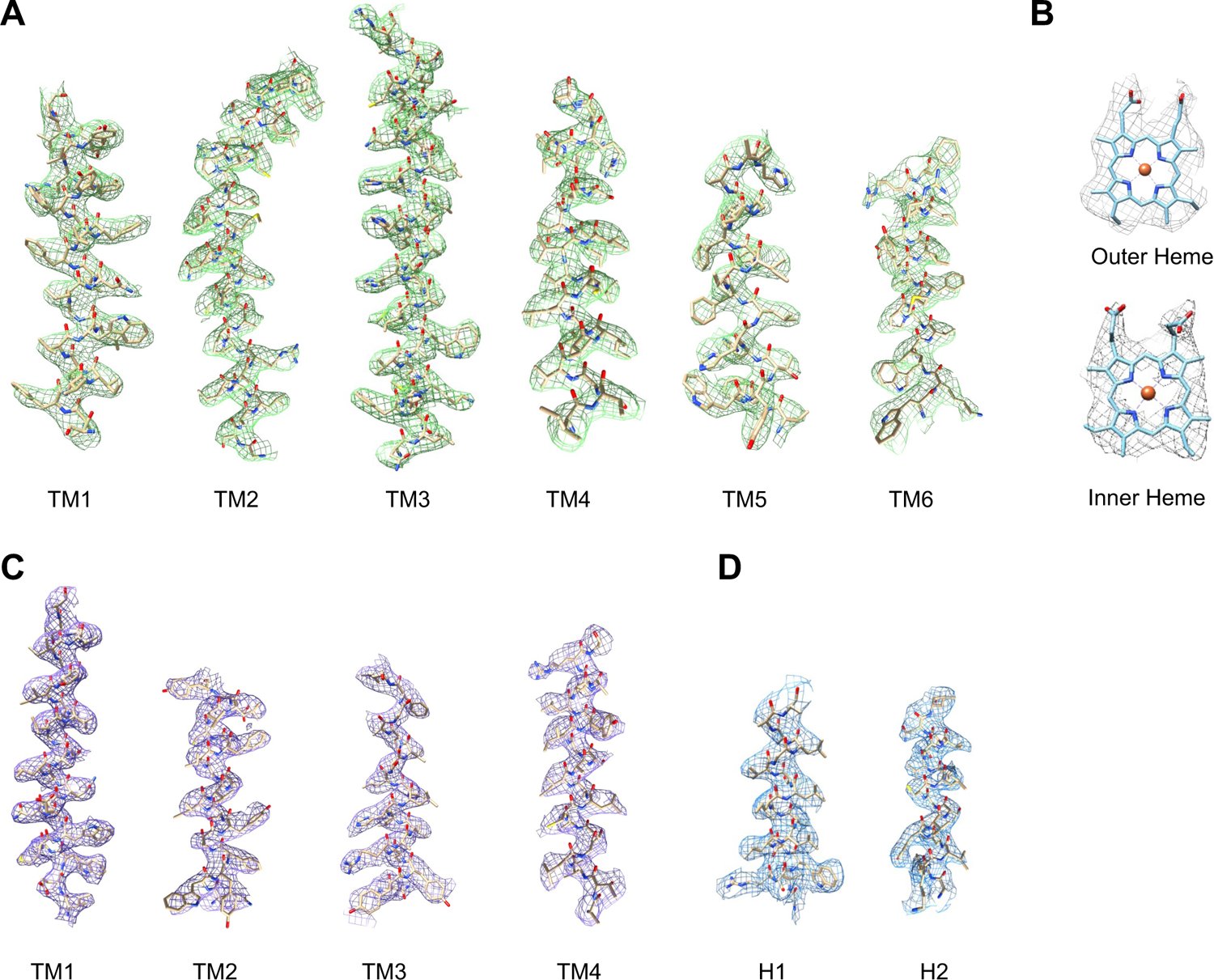
Representative density maps and models of NOX2, p22^phox^, and EROS. **A.** Representative electron density maps of NOX2 (TM1-TM6). **B.** Representative electron density maps of the two hemes. **C.** Representative electron density maps of p22^phox^ (TM1-TM4). **D.** Representative electron density maps of EROS (H1-H2).

**Figure S4.**
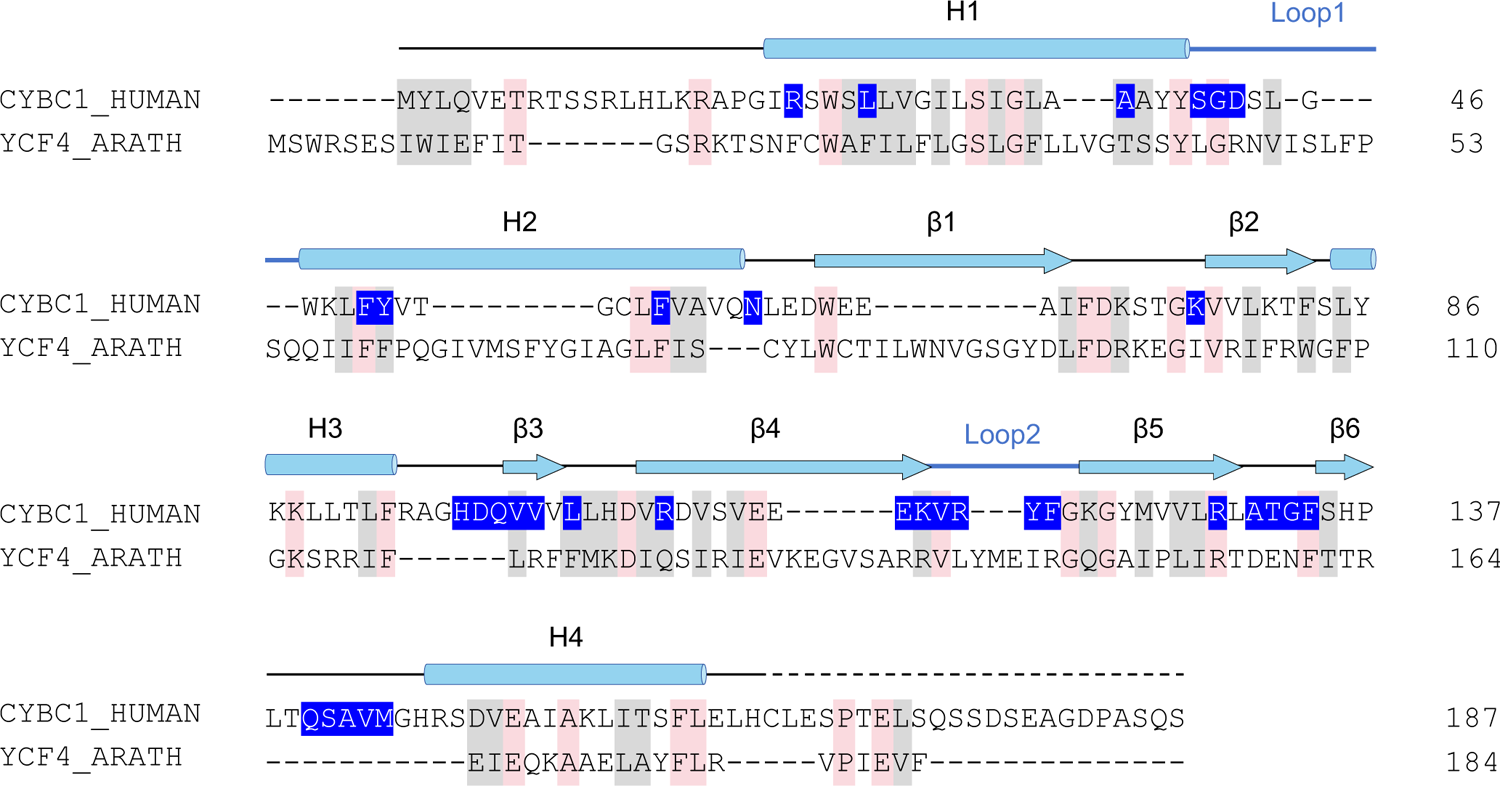
Sequence alignment of EROS and the *YCF4* gene product EROS is an ortholog of the plant protein encoded by *YCF4*, which is necessary for the expression of proteins belonging to the photosynthetic photosystem I complex. The *Homo sapiens* CYBC1 (UniProtKB: Q9BQA9) and *Arabidopsis thaliana* YCF4 (UniProtKB: P56788) sequences were aligned. Conserved residues are highlighted in pink and gray (pink: fully conserved, gray: highly conserved). Secondary structures are depicted as light blue cylinders (α helices), arrows (β sheets), and lines (loops). Dashed lines indicate unmodeled residues. The residues involved in NOX2 interactions are colored in blue.

**Table S1.** Cryo-EM data collection, model refinement and validation statistics.

| **Data collection and processing** | **NOX2-EROS** |
| --- | --- |
| Magnification | 105000 |
| Voltage (kV) | 300 |
| Electron exposure (e-/Å<sup>2</sup>) | 52.0 |
| Defocus range (µm) | -1.2~-2.5 |
| Pixel size (Å) | 0.85 |
| Symmetry imposed | C1 |
| Initial particle projections (no.) | 3470723 |
| Final particle projections (no.) | 131926 |
| **Refinement** |  |
| Initial model used (Alpha Fold database/PDB code) | 8GZ3 |
| Map resolution (Å) | 3.56 |
| FSC threshold | 0.143 |
| **Model composition** |  |
| Non-hydrogen atoms | 10209 |
| Protein residues | 1272 |
| **R.m.s. deviations** |  |
| Bond lengths (Å) | 0.006 |
| Bond angles (°) | 0.956 |
| **Validation** |  |
| MolProbity score | 1.85 |
| Clashes core | 8.43 |
| **Ramachandran plot** |  |
| Favored (%) | 94.19 |
| Allowed (%) | 5.81 |
| Outliers(%) | 0.00 |

